## Supplementary Figures for "Integrative study of skeletal muscle mitochondrial dysfunction in a murine pancreatic cancer-induced cachexia model"

**List of supplemental material:**

10 supplemental figures

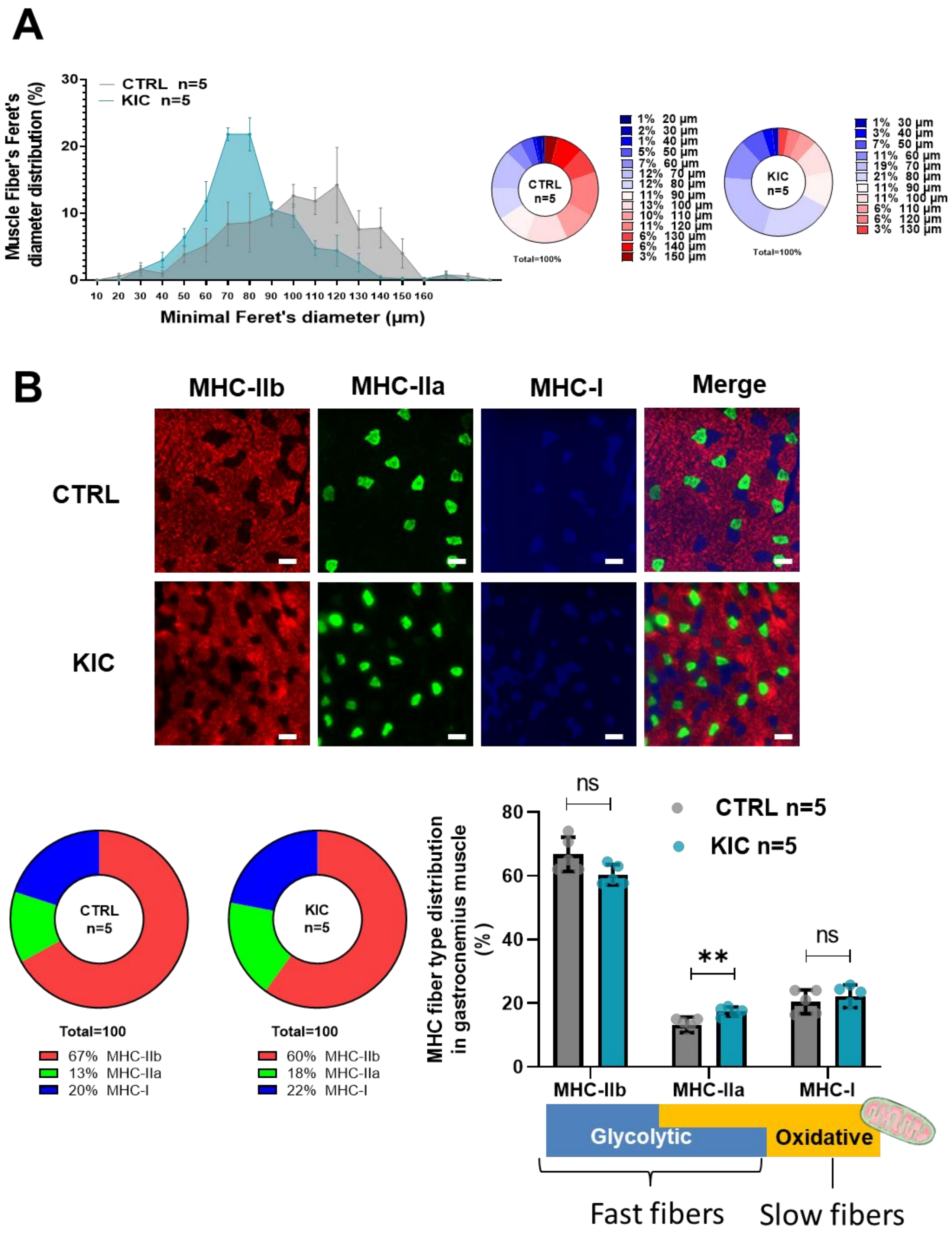
A
CTRL n=5
KIC n=5
Muscle Fiber's Feret's diameter distribution (%)
Minimal Feret's diameter (µm)
CTRL n=5
1% 20 µm
2% 30 µm
1% 40 µm
5% 50 µm
7% 60 µm
12% 70 µm
12% 80 µm
11% 90 µm
13% 100 µm
10% 110 µm
11% 120 µm
6% 130 µm
6% 140 µm
3% 150 µm
Total=100%
KIC n=5
1% 30 µm
3% 40 µm
7% 50 µm
11% 60 µm
19% 70 µm
21% 80 µm
11% 90 µm
11% 100 µm
6% 110 µm
6% 120 µm
3% 130 µm
Total=100%
B
MHC-IIb
MHC-IIa
MHC-I
Merge
CTRL
KIC
CTRL n=5
Total=100
67% MHC-IIb
13% MHC-IIa
20% MHC-I
KIC n=5
Total=100
60% MHC-IIb
18% MHC-IIa
22% MHC-I
MHC fiber type distribution in gastrocnemius muscle (%)
ns
**
ns
CTRL n=5
KIC n=5
MHC-IIb
MHC-IIa
MHC-I
Glycolytic
Oxidative
Fast fibers
Slow fibers

**Figure S1. Decreased fiber's size but moderate change in proportion of the different fiber types in PDAC atrophic muscle (related to Figure 2).**

**(A)** Muscle fiber's size distribution in gastrocnemius muscle from control and KIC mice. **(Left)** Minimal Feret's diameters were determined in gastrocnemius cross-sections from Laminin immunofluorescent staining (Figure 2B). Hundred fiber diameters were counted per group. The values are expressed as the percentage of the quantified fibers from 10 to 160 µm, and the image counts are representative of n=5 mice muscle cross sections used for each experimental condition. Values correspond to the mean ± SEM. The line represents the median distribution of minimal Feret diameters observed in healthy control (CTRL) versus cancer (KIC) mice. **(Right)** Mean percentage of muscle fiber's distribution for the control and cancer groups. **(B)** **(Top)** Representative immunostaining of transversal sections of gastrocnemius muscle from healthy control and cancer KIC male mice stained for MHC-IIb (red), MHC-IIa (green), and MHC-I (blue). Scale bars, 40µm. **(Bottom Left)** MHC-fiber-type distribution mean in gastrocnemius muscles from CTRL and KIC mice (n=5 mice/group). **(Bottom Right)** MHC-fiber-type distribution in gastrocnemius muscle from control and KIC mice (n=5 mice/group). Data are mean +/- SEM. Unpaired two-tailed Mann Whitney t-tests; ns = non-significant, **p<0.01. The scheme under the graph illustrates the different fiber type characteristics.

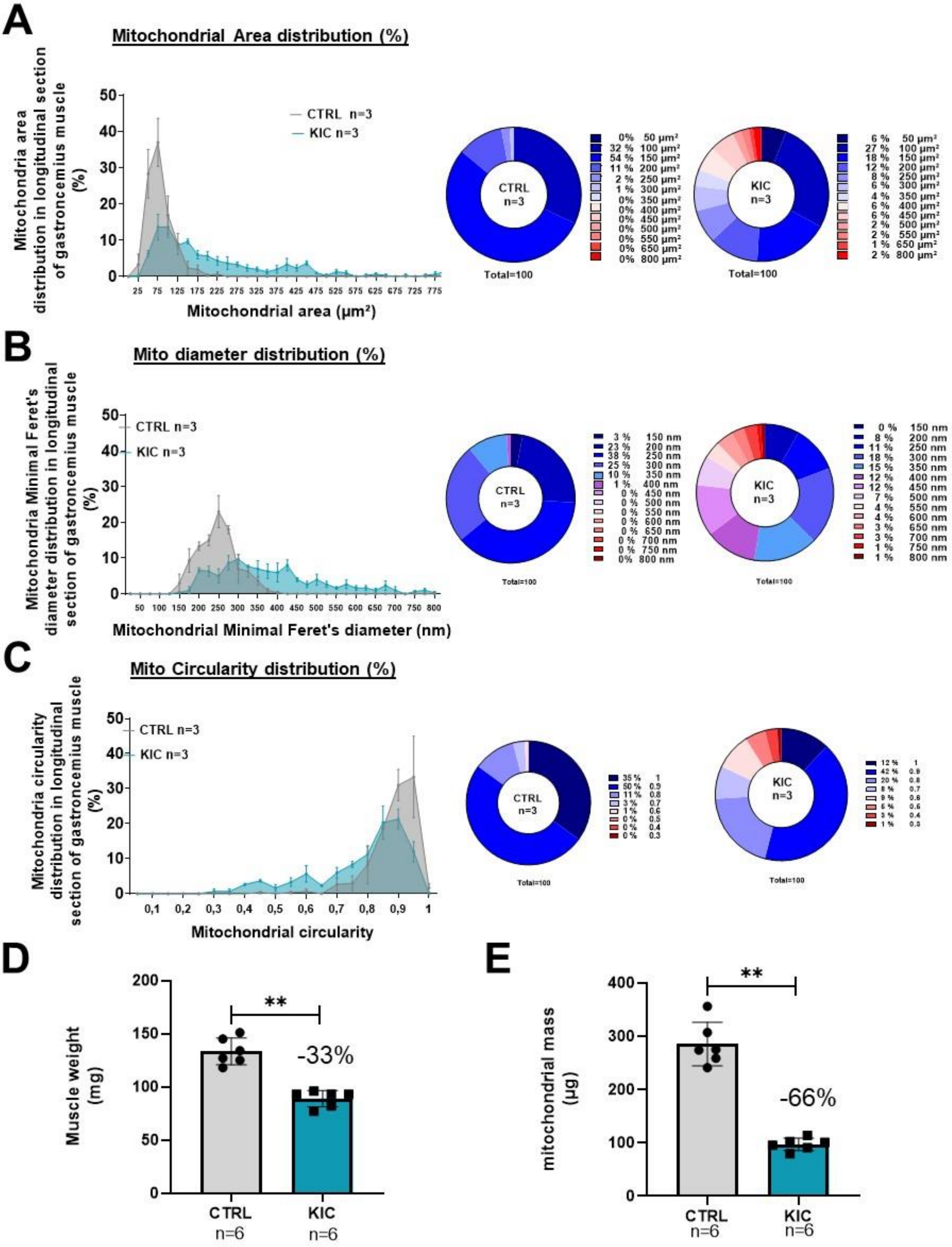
A
Mitochondrial Area distribution (%)
Mitochondria area distribution in longitudinal section of gastroncemius muscle (%)
CTRL n=3
KIC n=3
Mitochondrial area (µm²)
CTRL n=3
0% 50 µm²
32 % 100 µm²
54 % 150 µm²
11 % 200 µm²
2 % 250 µm²
1 % 300 µm²
0% 350 µm²
0% 400 µm²
0% 450 µm²
0% 500 µm²
0% 550 µm²
0% 650 µm²
0% 800 µm²
Total=100
KIC n=3
6 % 50 µm²
27 % 100 µm²
18 % 150 µm²
12 % 200 µm²
8 % 250 µm²
6 % 300 µm²
4 % 350 µm²
6 % 400 µm²
6 % 450 µm²
2 % 500 µm²
2 % 550 µm²
1 % 650 µm²
2 % 800 µm²
Total=100
B
Mito diameter distribution (%)
Mitochondria Minimal Feret's diameter distribution in longitudinal section of gastroncemius muscle (%)
CTRL n=3
KIC n=3
Mitochondrial Minimal Feret's diameter (nm)
CTRL n=3
3 % 150 nm
23 % 200 nm
38 % 250 nm
25 % 300 nm
10 % 350 nm
1 % 400 nm
0 % 450 nm
0 % 500 nm
0 % 550 nm
0 % 600 nm
0 % 650 nm
0 % 700 nm
0 % 750 nm
0% 800 nm
Total=100
KIC n=3
0 % 150 nm
8 % 200 nm
11 % 250 nm
18 % 300 nm
15 % 350 nm
12 % 400 nm
12 % 450 nm
7 % 500 nm
4 % 550 nm
4 % 600 nm
3 % 650 nm
3 % 700 nm
1 % 750 nm
1 % 800 nm
Total=100
C
Mito Circularity distribution (%)
Mitochondria circularity distribution in longitudinal section of gastroncemius muscle (%)
CTRL n=3
KIC n=3
Mitochondrial circularity
CTRL n=3
35 % 1
50 % 0.9
11 % 0.8
3 % 0.7
1 % 0.6
0 % 0.5
0 % 0.4
0 % 0.3
Total=100
KIC n=3
12 % 1
42 %
20 %
Total=100
D
Muscle weight (mg)
**
-33%
CTRL n=6
KIC n=6
E
mitochondrial mass (µg)
**
-66%
CTRL n=6
KIC n=6

**Figure S2. Mitochondrial morphology is profoundly altered in KIC muscles (related to Figure 5).**

**(A-C)** Percentages (%) for mitochondria morphological characteristics in longitudinal sections of gastrocnemius muscles from healthy CTRL and cancer KIC mice, analyzed by TEM. **(A)** mitochondrial area ($\mu m^2$), **(B)** mitochondrial minimal Feret's diameter, and **(C)** mitochondrial circularity. n=3 mice/group and 100 mitochondria measured/mouse. **(D)** Muscle weight of entire gastrocnemius muscle used for mitochondria isolation shown in Figure S2E, and to normalize mitochondrial mass quantity shown in Figure 5D. Data are mean +/- SD; n=6 mice/group. Unpaired two-tailed Mann Whitney t-tests; **p<0.01. **(E)** Total mass (in protein equivalent) of mitochondria isolated from entire gastrocnemius muscle from CTRL and KIC mice. Data are mean +/- SD; n=6 mice/group. Unpaired two-tailed Mann Whitney t-tests; **p<0.01.

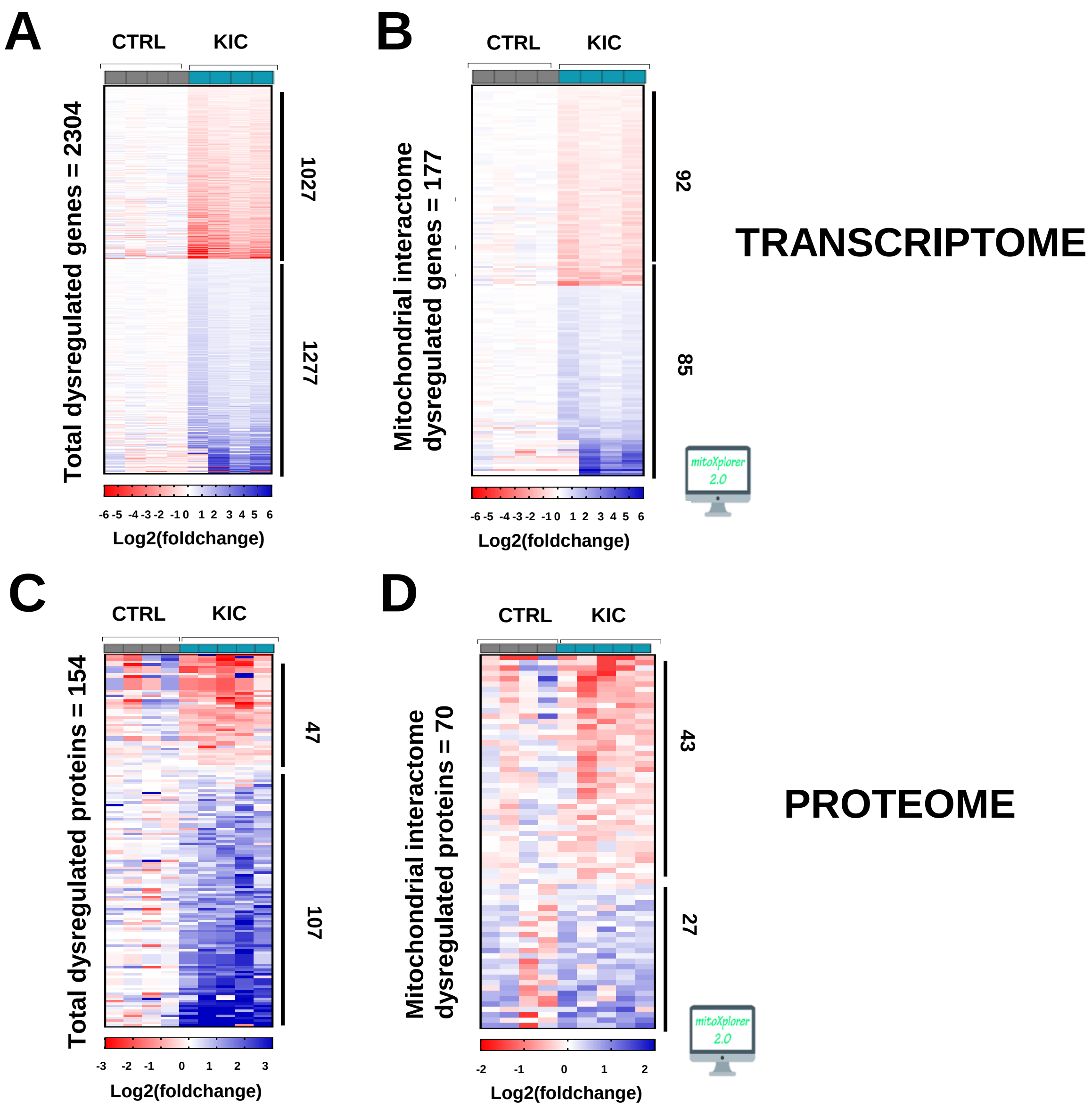


**Figure S3. Transcriptomic and proteomic analysis of cancer sarcopenic muscles.**
**(A)** Heatmap representing total number of dysregulated genes (FDR<0.05) in gastrocnemius muscle from cancer KIC mice (n=4) compared to healthy CTRL mice (n=4). **(B)** Heatmap representing dysregulated genes from mitochondrial interactome after MitoXplorer selection (FDR<0.05) in gastrocnemius muscle from the same mice as (A). **(C)** Heatmap representing total number of dysregulated proteins in gastrocnemius muscle from cancer KIC mice (n=5) compared to healthy CTRL mice (n=4). **(D)** Heatmap representing dysregulated proteins from mitochondrial interactome after MitoXplorer selection in gastrocnemius muscle from the same mice as (C).

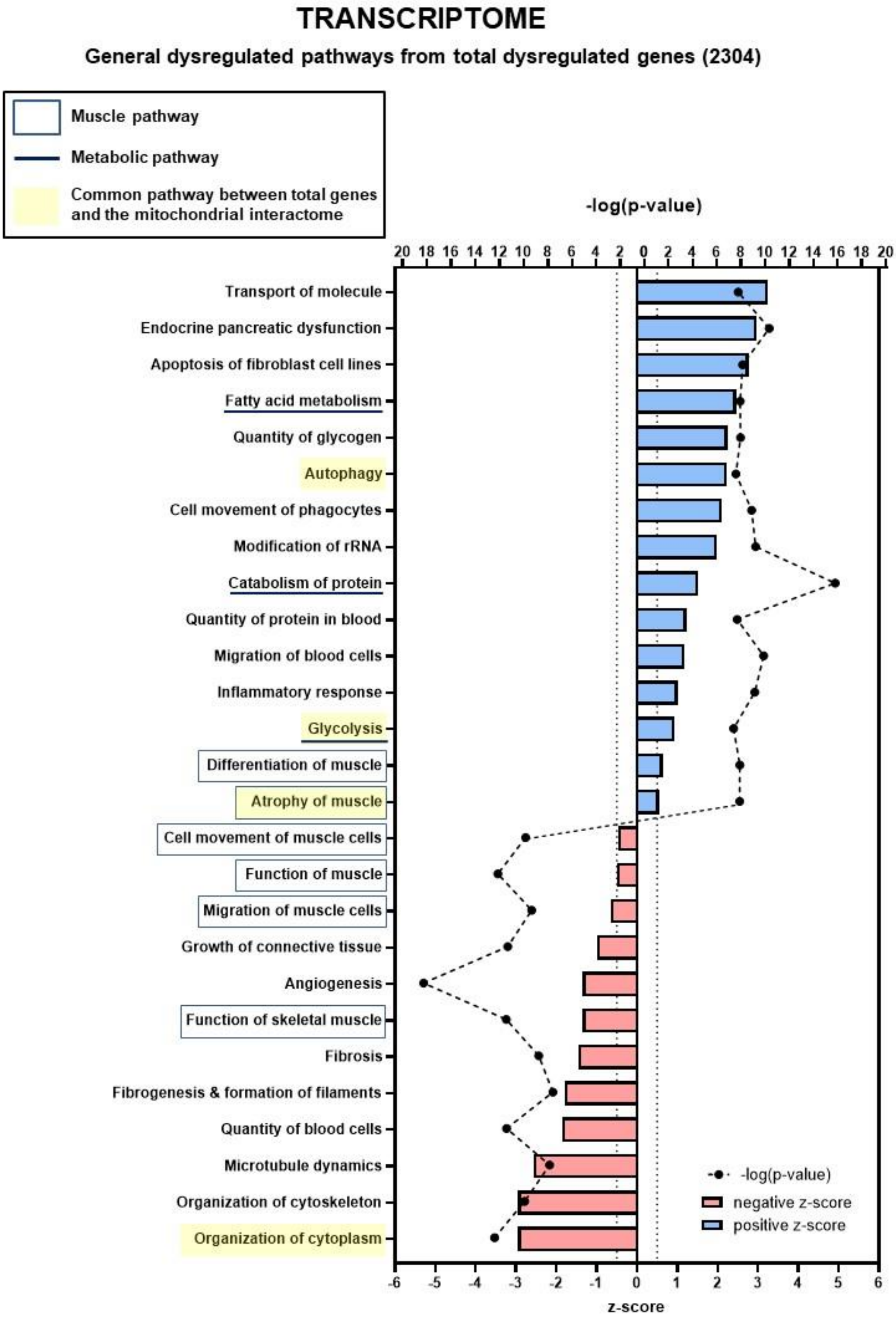


**Figure S4. Transcriptomic analysis of total dysregulated genes in cancer sarcopenic muscles.** Ingenuity Pathways Analysis (IPA) showing functions, pathways and diseases significantly dysregulated in gastrocnemius muscle of cancer KIC mice compared to healthy control mice, associated with all genes with significant dysregulation (FDR>0.05) = 2304 genes. Each process is represented using z-score>0.5 as upregulated (blue) or downregulated (red), and the significance represented as -log(p-value) with black dots connected with a dotted line.

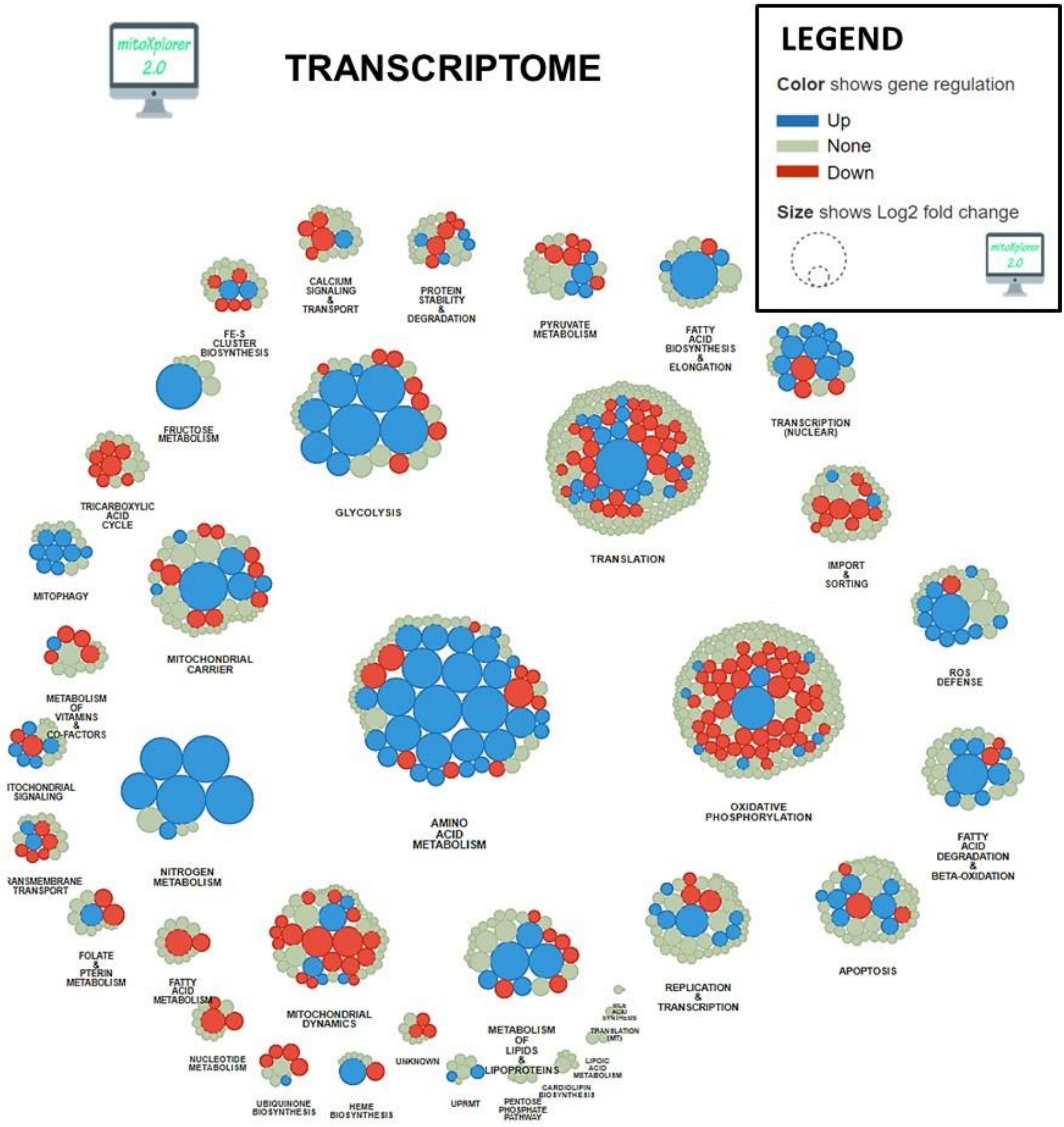


**Figure S5.** Mitochondrial interactome of down- or up-regulated genes involved in mitochondrial processes in KIC gastrocnemius muscle compared to CTRL (muscle RNA n=4 for each condition). Genes have been selected from total dysregulated genes with FDR<0.05, and affiliated to processes after enrichment using MitoXplorer tool.

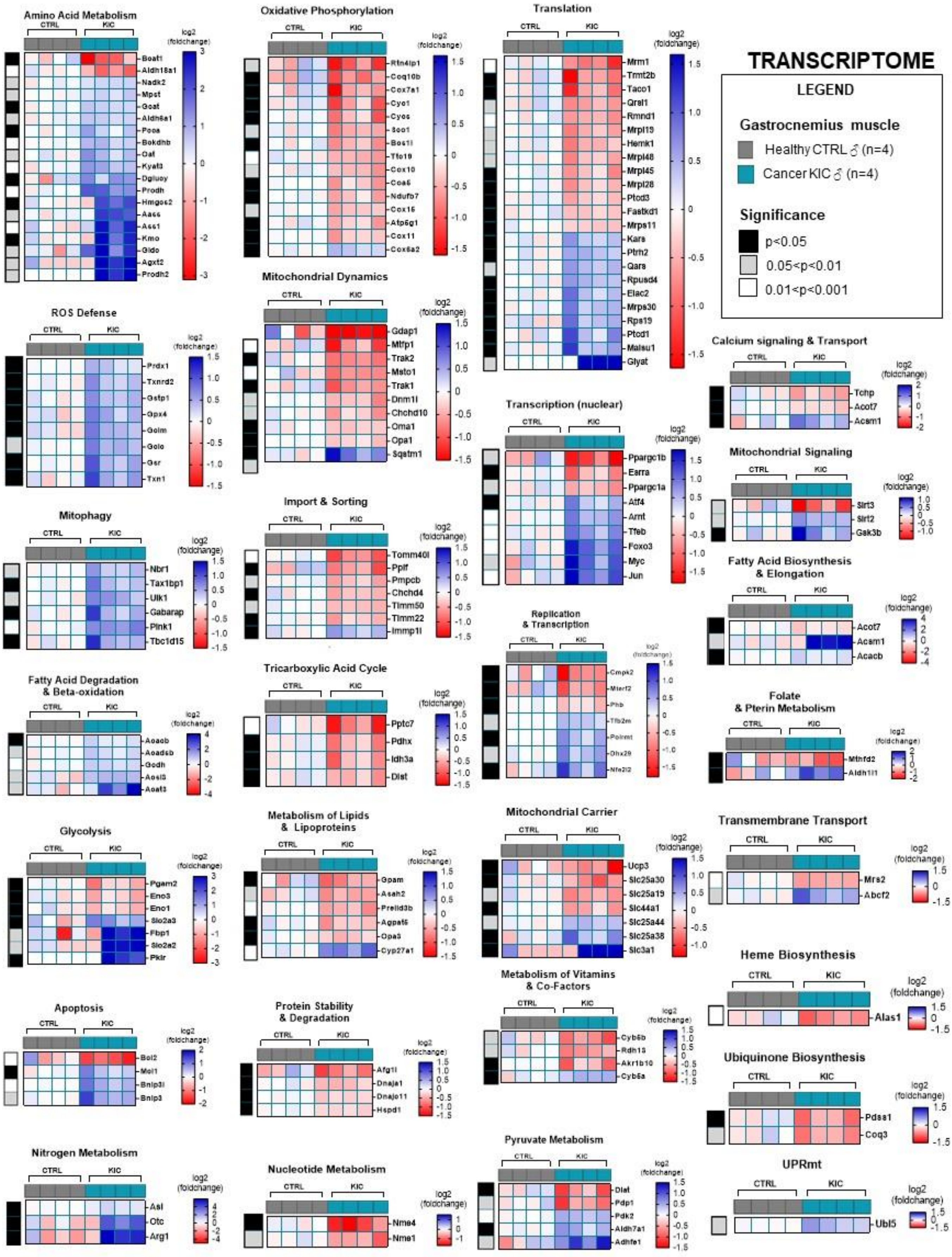
TRANSCRIPTOME
LEGEND
Gastrocnemius muscle
Healthy CTRL ♂ (n=4)
Cancer KIC ♂ (n=4)
Significance
p<0.05
0.05<p<0.01
0.01<p<0.001
CTRL
KIC
log2 (foldchange)
Amino Acid Metabolism
Bcat1
Aldh18a1
Nadk2
Mpst
Goat
Aldh6a1
Pcca
Bckdhb
Oat
Kyat3
Dglucy
Prodh
Hmgcs2
Aass
Ass1
Kmo
Gldc
Agxt2
Prodh2
Oxidative Phosphorylation
Rtn4ip1
Coq10b
Cox7a1
Cyc1
Cycs
Sco1
Bcs1l
Tfc19
Cox10
Coa6
Ndufb7
Cox15
Atp5g1
Cox11
Cox6a2
Translation
Mrm1
Trmt2b
Taco1
Qrsl1
Rmnd1
Mrpl19
Hemk1
Mrpl48
Mrpl45
Mrpl28
Ptcd3
Fastkd1
Mrps11
Kars
Ptrh2
Qars
Rpusd4
Elac2
Mrps30
Rps19
Ptcd1
Malsu1
Glyat
ROS Defense
Prdx1
Txnrd2
Gstp1
Gpx4
Gclm
Gclc
Gsr
Txn1
Mitochondrial Dynamics
Gdap1
Mtfp1
Trak2
Msto1
Trak1
Dnm1l
Chchd10
Oma1
Opa1
Sqstm1
Transcription (nuclear)
Ppargc1b
Esrra
Ppargc1a
Atf4
Arnt
Tfeb
Foxo3
Myc
Jun
Calcium signaling & Transport
Tchp
Acot7
Acsm1
Mitochondrial Signaling
Sirt3
Sirt2
Gsk3b
Mitophagy
Nbr1
Tax1bp1
Ulk1
Gabarap
Pink1
Tbc1d15
Import & Sorting
Tomm40l
Ppif
Pmpcb
Chchd4
Timm50
Timm22
Immp1l
Replication & Transcription
Fatty Acid Biosynthesis & Elongation
Acot7
Acsm1
Acacb
Fatty Acid Degradation & Beta-oxidation
Acadl
Acadsb
Gcdh
Acsl3
Acat3
Tricarboxylic Acid Cycle
Pptc7
Pdhx
Idh3a
Dlst
Folate & Pterin Metabolism
Mthfd2
Aldh1l1
Glycolysis
Pgam2
Eno3
Eno1
Slc2a3
Fbp1
Slc2a2
Pklr
Metabolism of Lipids & Lipoproteins
Gpam
Asah2
Prelid3b
Agpat5
Cpa3
Cyp27a1
Mitochondrial Carrier
Ucp3
Slc25a30
Slc25a19
Slc44a1
Slc25a44
Slc25a38
Slc3a1
Transmembrane Transport
Mrs2
Abcf2
Heme Biosynthesis
Alas1
Metabolism of Vitamins & Co-Factors
Cyb5b
Rdh13
Akr1b10
Cyb5a
Apoptosis
Bcl2
Mcl1
Bnip3l
Bnip3
Protein Stability & Degradation
Afg1l
Dnaja1
Dnajc11
Hspd1
Ubiquinone Biosynthesis
Pdss1
Coq3
Nitrogen Metabolism
Asl
Otc
Arg1
Nucleotide Metabolism
Nme4
Nme1
Pyruvate Metabolism
Dlat
Pdp1
Pdk2
Aldh7a1
Adhfe1
UPRmt
Ubl5

**Figure S6. Cancer sarcopenic muscles demonstrate obvious mitochondria-associated genes dysregulation (related to Figure S5).**

Heatmaps showing log2 (Fold change) of differentially dysregulated genes involved in mitochondrial processes (based on FDR<0.05) in cancer KIC gastrocnemius muscle compared to healthy CTRL (muscle RNA n=4 for each condition).

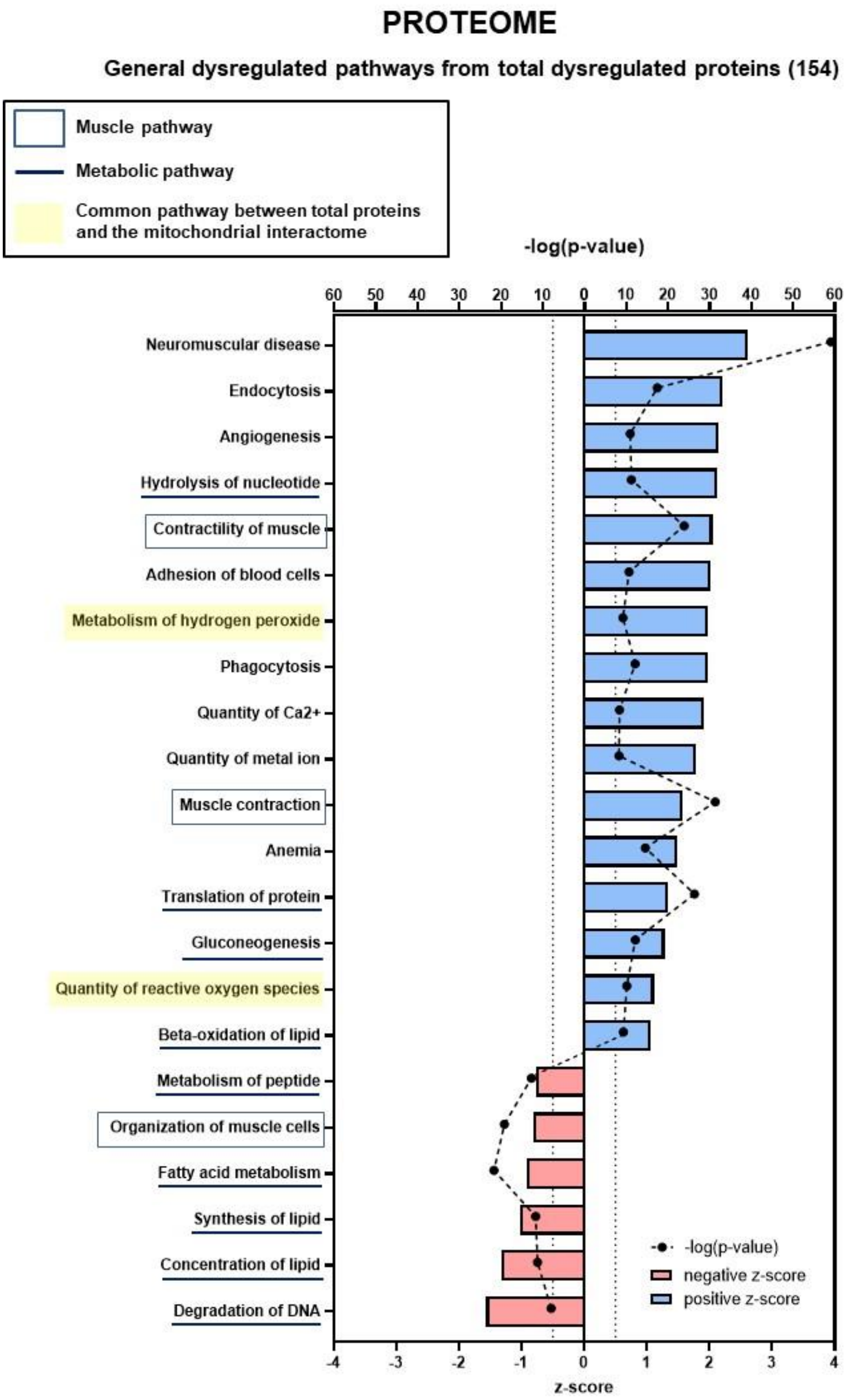


**Figure S7. Proteomic analysis showing total proteins dysregulated in cancer sarcopenic muscle.**

IPA enrichment analysis of dysregulated functions, pathways and diseases in gastrocnemius muscle of cancer KIC compared to control mice, associated with all proteins with significant dysregulations (p-value<0.05) = 360 proteins. Each process is represented using z-score>0.5 as upregulated (blue) or downregulated (red), and the significance represented as –log(p-value) with black dots connected with a dotted line.

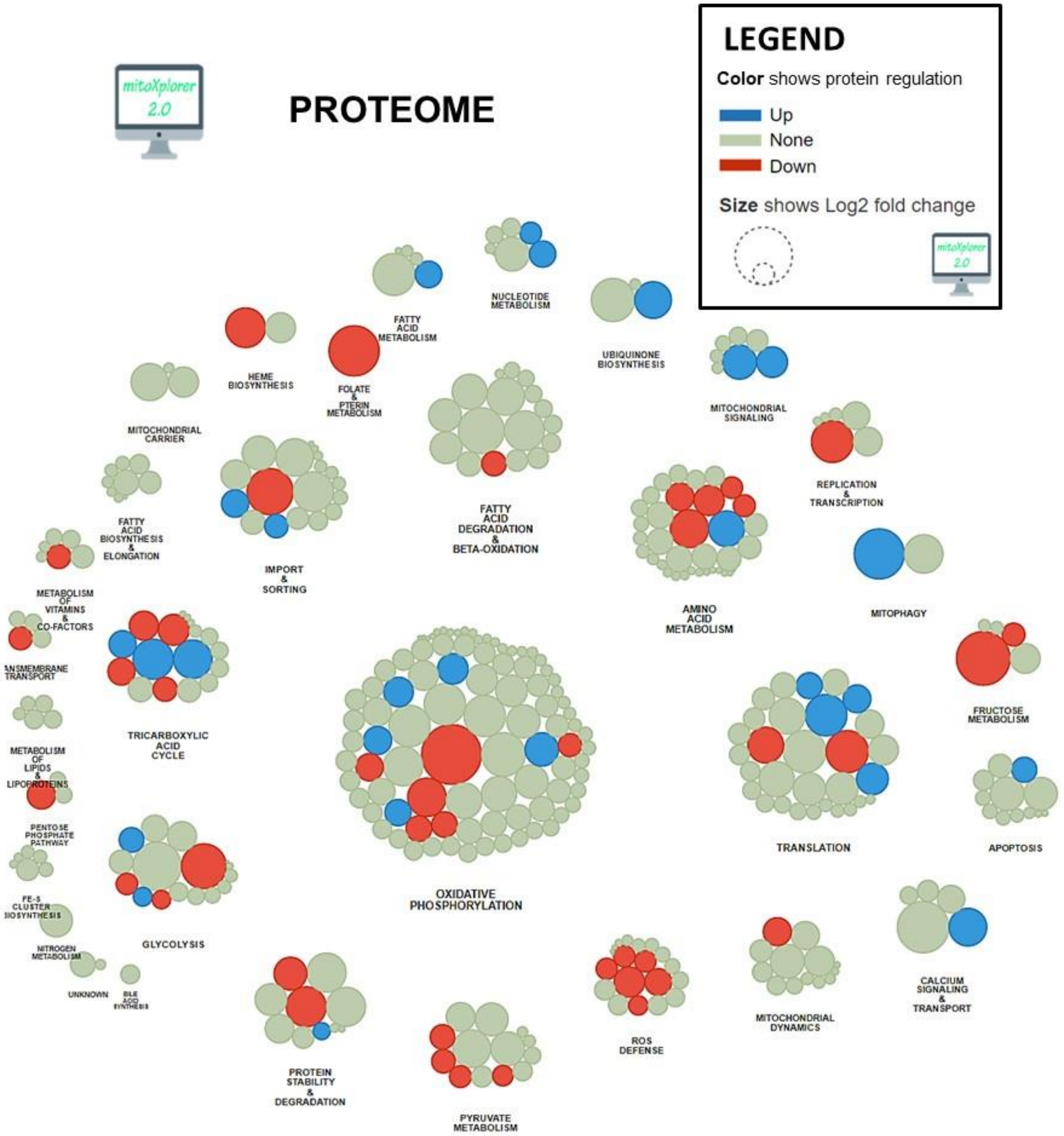


**Figure S8.** Mitochondrial interactome of down- or up-regulated proteins involved in mitochondrial processes in KIC gastrocnemius muscle compared to CTRL. Proteins have been selected from total dysregulated proteins and affiliated to processes after enrichment using MitoXplorer tool.

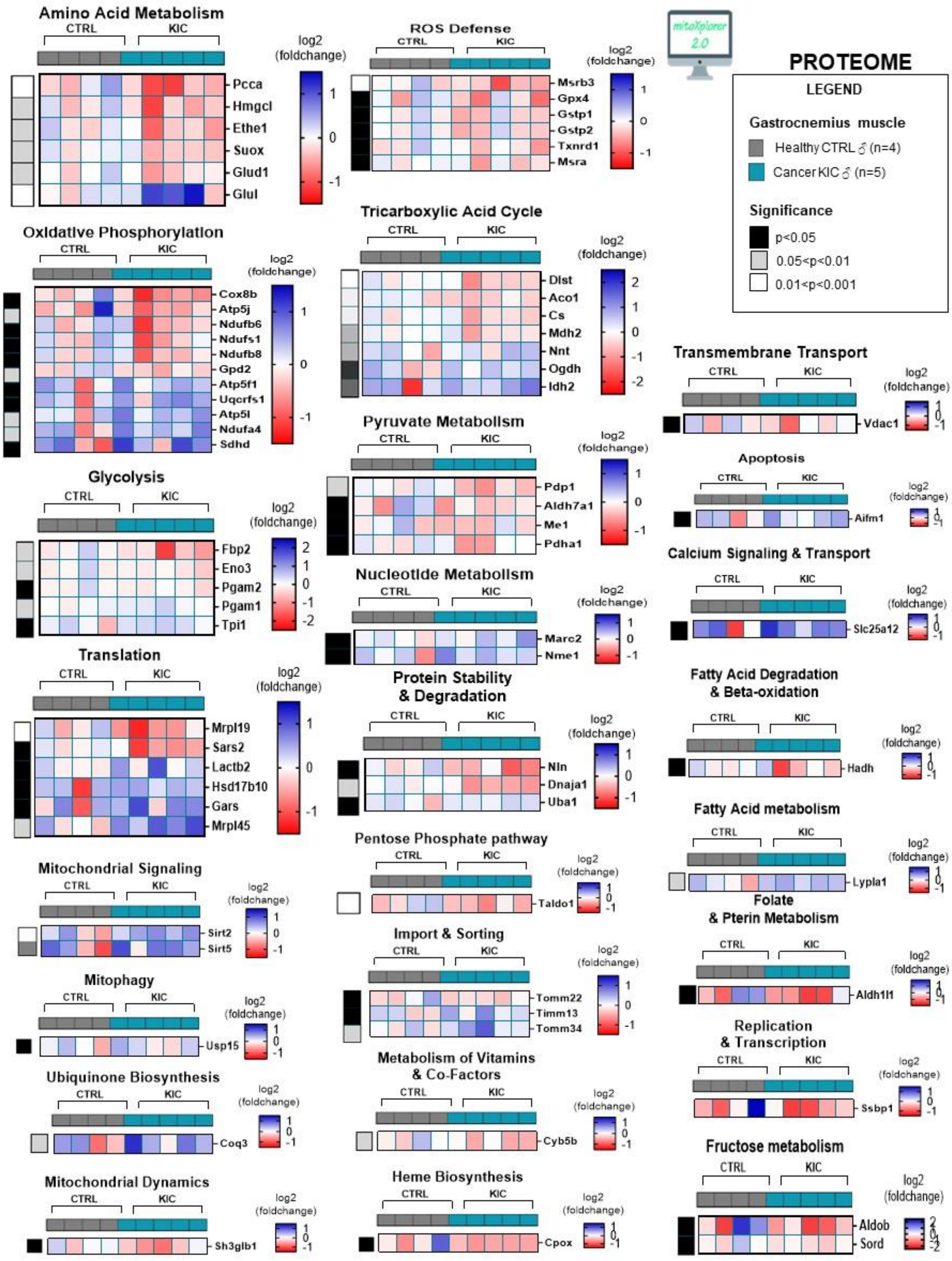
PROTEOME
mitoXplorer 2.0
LEGEND
Gastrocnemius muscle
Healthy CTRL ♂ (n=4)
Cancer KIC ♂ (n=5)
Significance
p<0.05
0.05<p<0.01
0.01<p<0.001
Amino Acid Metabolism
CTRL
KIC
Pcca
Hmgcl
Ethe1
Suox
Glud1
Glul
log2 (foldchange)
ROS Defense
Msrb3
Gpx4
Gstp1
Gstp2
Txnrd1
Msra
Oxidative Phosphorylation
Cox8b
Atp5j
Ndufb6
Ndufs1
Ndufb8
Gpd2
Atp5f1
Uqcrfs1
Atp5l
Ndufa4
Sdhd
Tricarboxylic Acid Cycle
Dlst
Aco1
Cs
Mdh2
Nnt
Ogdh
Idh2
Transmembrane Transport
Vdac1
Glycolysis
Fbp2
Eno3
Pgam2
Pgam1
Tpi1
Pyruvate Metabolism
Pdp1
Aldh7a1
Me1
Pdha1
Apoptosis
Aifm1
Calcium Signaling & Transport
Slc25a12
Nucleotide Metabolism
Marc2
Nme1
Translation
Mrpl19
Sars2
Lactb2
Hsd17b10
Gars
Mrpl45
Protein Stability & Degradation
Nln
Dnaja1
Uba1
Fatty Acid Degradation & Beta-oxidation
Hadh
Fatty Acid metabolism
Lypla1
Mitochondrial Signaling
Sirt2
Sirt5
Pentose Phosphate pathway
Taldo1
Folate & Pterin Metabolism
Aldh1l1
Mitophagy
Usp15
Import & Sorting
Tomm22
Timm13
Tomm34
Replication & Transcription
Ssbp1
Ubiquinone Biosynthesis
Coq3
Metabolism of Vitamins & Co-Factors
Cyb5b
Fructose metabolism
Aldob
Sord
Mitochondrial Dynamics
Sh3glb1
Heme Biosynthesis
Cpox

**Figure S9. Cancer sarcopenic muscles demonstrate obvious mitochondria-associated protein dysregulation (related to Figure S8).**

Heatmap representing dysregulated proteins from mitochondrial interactome after MitoXplorer selection (p-value<0,05, peptide>1, log2(Foldchange)>0,5) in same gastrocnemius muscles as in Figure S8.

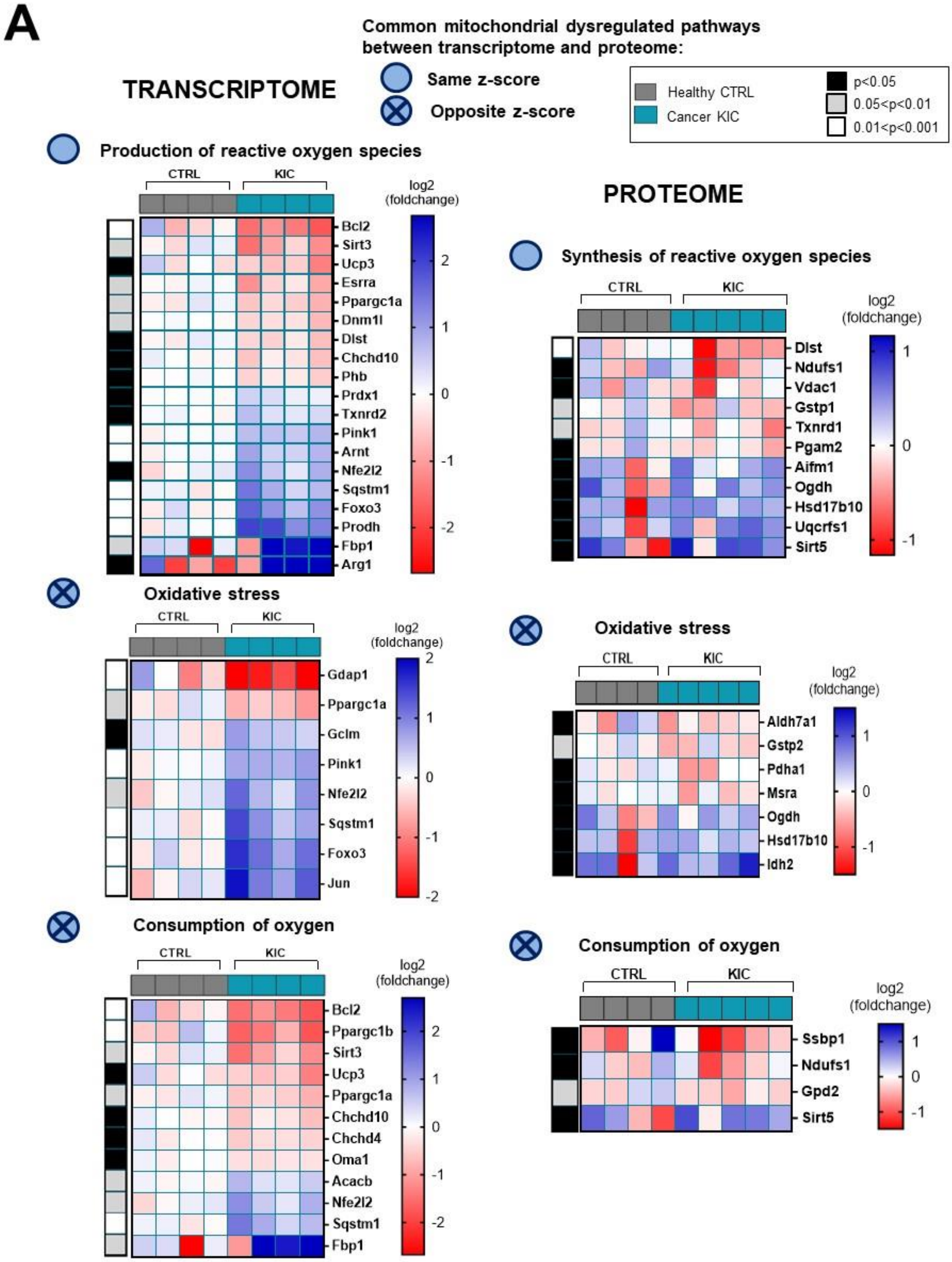
A
Common mitochondrial dysregulated pathways between transcriptome and proteome:
Same z-score
Opposite z-score
Healthy CTRL
Cancer KIC
p<0.05
0.05<p<0.01
0.01<p<0.001
TRANSCRIPTOME
Production of reactive oxygen species
CTRL
KIC
log2 (foldchange)
Bcl2
Sirt3
Ucp3
Esrra
Ppargc1a
Dnm1l
Dlst
Chchd10
Phb
Prdx1
Txnrd2
Pink1
Arnt
Nfe2l2
Sqstm1
Foxo3
Prodh
Fbp1
Arg1
Oxidative stress
Gdap1
Ppargc1a
Gclm
Pink1
Nfe2l2
Sqstm1
Foxo3
Jun
Consumption of oxygen
Bcl2
Ppargc1b
Sirt3
Ucp3
Ppargc1a
Chchd10
Chchd4
Oma1
Acacb
Nfe2l2
Sqstm1
Fbp1
PROTEOME
Synthesis of reactive oxygen species
Dlst
Ndufs1
Vdac1
Gstp1
Txnrd1
Pgam2
Aifm1
Ogdh
Hsd17b10
Uqcrfs1
Sirt5
Oxidative stress
Aldh7a1
Gstp2
Pdha1
Msra
Ogdh
Hsd17b10
Idh2
Consumption of oxygen
Ssbp1
Ndufs1
Gpd2
Sirt5

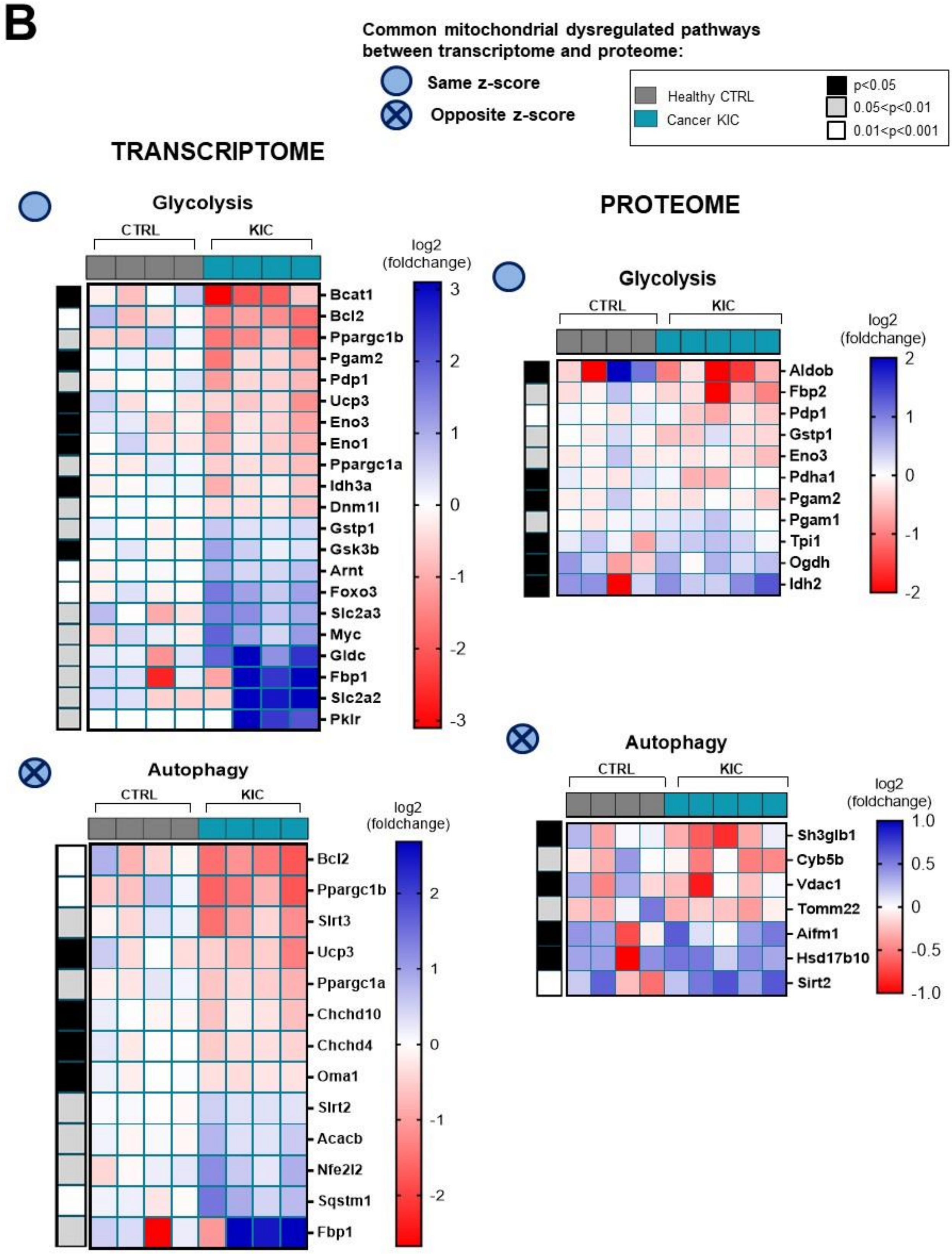
B
Common mitochondrial dysregulated pathways between transcriptome and proteome:
Same z-score
Opposite z-score
Healthy CTRL
Cancer KIC
p<0.05
0.05<p<0.01
0.01<p<0.001
TRANSCRIPTOME
Glycolysis
CTRL
KIC
log2 (foldchange)
Bcat1
Bcl2
Ppargc1b
Pgam2
Pdp1
Ucp3
Eno3
Eno1
Ppargc1a
Idh3a
Dnm1l
Gstp1
Gsk3b
Arnt
Foxo3
Slc2a3
Myc
Gldc
Fbp1
Slc2a2
Pklr
3
2
1
0
-1
-2
-3
Autophagy
CTRL
KIC
log2 (foldchange)
Bcl2
Ppargc1b
Sirt3
Ucp3
Ppargc1a
Chchd10
Chchd4
Oma1
Sirt2
Acacb
Nfe2l2
Sqstm1
Fbp1
2
1
0
-1
-2
PROTEOME
Glycolysis
CTRL
KIC
log2 (foldchange)
Aldob
Fbp2
Pdp1
Gstp1
Eno3
Pdha1
Pgam2
Pgam1
Tpi1
Ogdh
Idh2
2
1
0
-1
-2
Autophagy
CTRL
KIC
log2 (foldchange)
Sh3glb1
Cyb5b
Vdac1
Tomm22
Aifm1
Hsd17b10
Sirt2
1.0
0.5
0
-0.5
-1.0

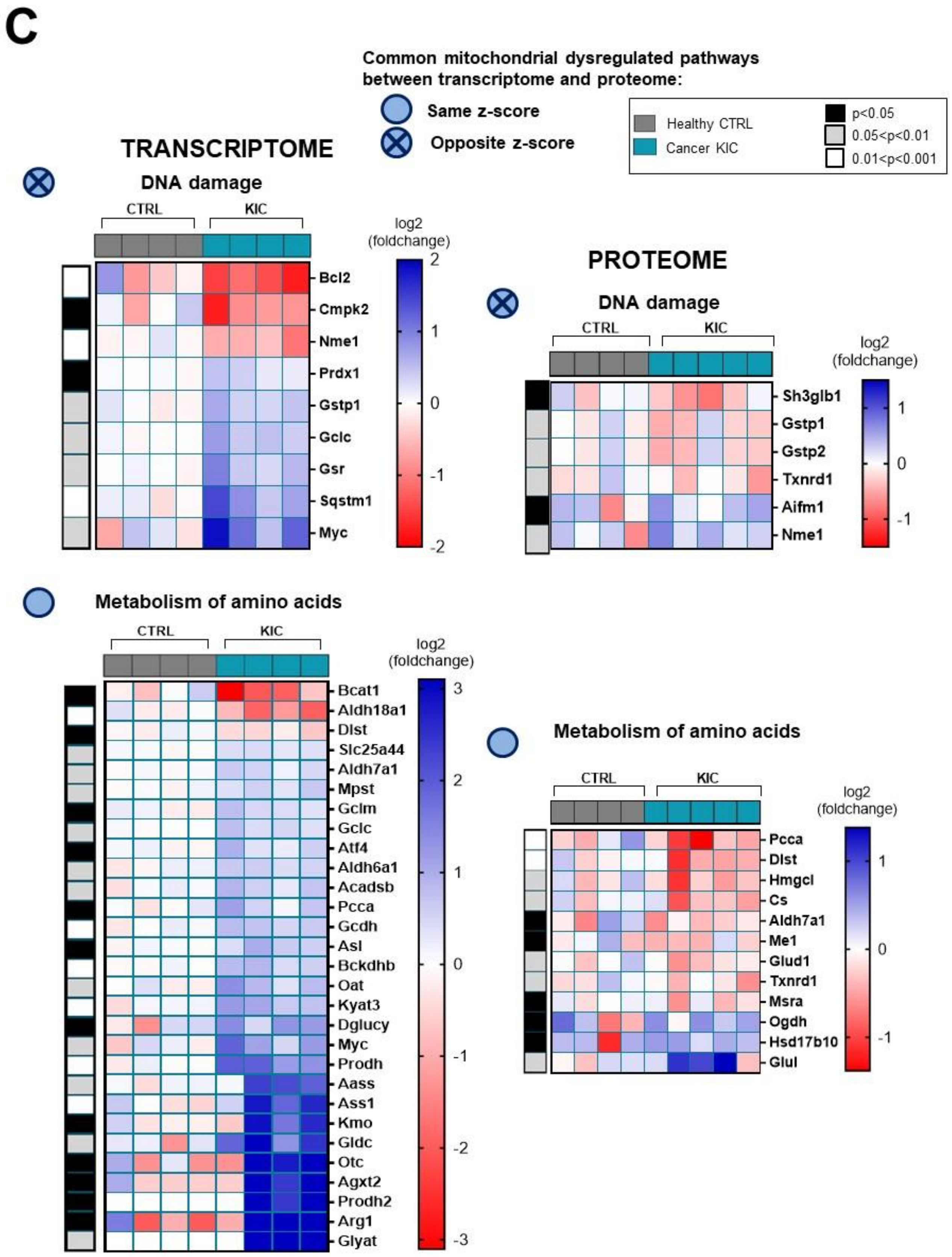
C
Common mitochondrial dysregulated pathways between transcriptome and proteome:
Same z-score
Opposite z-score
Healthy CTRL
Cancer KIC
p<0.05
0.05<p<0.01
0.01<p<0.001
TRANSCRIPTOME
DNA damage
CTRL
KIC
log2 (foldchange)
Bcl2
Cmpk2
Nme1
Prdx1
Gstp1
Gclc
Gsr
Sqstm1
Myc
PROTEOME
DNA damage
Sh3glb1
Gstp1
Gstp2
Txnrd1
Aifm1
Nme1
Metabolism of amino acids
Bcat1
Aldh18a1
Dlst
Slc25a44
Aldh7a1
Mpst
Gclm
Gclc
Atf4
Aldh6a1
Acadsb
Pcca
Gcdh
Asl
Bckdhb
Oat
Kyat3
Dglucy
Myc
Prodh
Aass
Ass1
Kmo
Gldc
Otc
Agxt2
Prodh2
Arg1
Glyat
Metabolism of amino acids
Pcca
Dlst
Hmgcl
Cs
Aldh7a1
Me1
Glud1
Txnrd1
Msra
Ogdh
Hsd17b10
Glul

**Figures S10. Common mitochondrial dysregulated pathways between transcriptomic and proteomic analyses (related to Figures 6 and 7)**
**(A-C)** Heatmaps corresponding to differential pathways in transcriptome (left) and proteome (right) analyses, to compare side-by-side the pathways that are similarly affected at mRNA and protein levels.
